# Quantitative analysis of transporter activity biosensors

**DOI:** 10.1101/2020.09.03.282301

**Authors:** Jihyun Park, Taylor M. Chavez, Wolf B. Frommer, Lily S. Cheung

**Affiliations:** School of Chemical and Biomolecular Engineering. Georgia Institute of Technology. Atlanta, GA 30332, USA; Department of Bioengineering. Stanford University. Stanford, CA 94305, USA; Institute for Molecular Physiology and Cluster of Excellence on Plant Sciences (CEPLAS), Heinrich Heine Universität Düsseldorf; Institute of Transformative Bio-Molecules (WPI-ITbM), Nagoya University, Aichi, Japan

**Keywords:** Biomolecular sensors, SWEET sugar transporters, fluorescence, genetically encoded, activity sensor

## Abstract

The allocation of sugars from photosynthetic leaves to storage tissues in seeds, fruits, and tubers is an important determinant of crop yields. Genetically guided selection and transgenic modification of plant membrane transporters can help enhance crop yields and increase pathogen resistance. Yet, quantitative, systems-level models to support this effort are lacking. Recently, biosensors gained popularity for collecting spatiotemporally resolved information on cell physiology and validating computational models. Here, we report the design and use of genetically-encoded biosensors to measure the activity of SWEETs, the only family of sugar transporters known to facilitate the cellular release of sugar in plants. We created SweetTrac sensors by inserting circularly-permutated GFP into SWEET transporters, resulting in chimeras that translate substrate-triggered conformational rearrangements during the transport cycle into detectable changes in fluorescence intensity. We demonstrate that a combination of cell sorting and bioinformatics can be applied as a general approach to accelerate the design of biosensors for *in vivo* biochemistry. Finally, mass action kinetics analysis of the biosensors’ response suggests that SWEETs are low-affinity, near-symmetric transporters that can rapidly equilibrate intra- and extracellular concentrations of sugars.

**Significance Statement:** Transporters are the gatekeepers of the cell. Transporters facilitate the exchange of ions and metabolites between cellular and subcellular compartments, thus controlling processes from bacterial chemotaxis to the release of neurotransmitters. In plants, transporters play critical roles in the allocation of carbon to different organs. Biosensors derived from transporters have been generated to monitor the activity of these proteins within the complex environment of the cell. However, a quantitative framework that reconciles molecular and cellular-level events to help interpret the response of biosensors is still lacking. Here, we created novel sugar transport biosensors and formulated a mathematical model to explain their response. These types of models can help realize multiscale, dynamic simulations of metabolite allocation to guide crop improvement.

## Introduction

Membrane transporters are key regulators of metabolism. In plants, sugar transporters are responsible for controlling the long-distance translocation of sugars from photosynthetic tissues to sinks like roots, seeds, and fruits. The cellular uptake and release of sugars is one of the most intensely studied processes in biochemistry and plant physiology (1).

Despite their recognized importance, determining the turnover numbers and affinities of sugar transporters remains difficult. In general, biochemical characterization of transporters is challenging because, unlike soluble proteins, they must be embedded in a membrane to work. Transporter activity is usually measured using radiotracers, microelectrodes, or substrate analogs in heterologous systems like yeast and *Xenopus* oocytes (2). Physiological studies of transporters *in planta* are also challenging since mutant phenotypes are pleiotropic, and paralogs commonly display functional redundancy. As a result, only a small fraction of plant transporters have been characterized. For example, the function of less than a quarter of the ~1,200 transporters in the model plant *Arabidopsis thaliana* have been validated experimentally (3). In contrast, only 2.3% of the ~1400 currently available *Arabidopsis* structures in the Protein Data Bank are transporters.

New technologies to characterize transporters in their natural environment are necessary to understand their contribution to plant physiology fully. The expression of tagged proteins in plants helps ascertain localization and even protein levels, but they do not show whether the transporter is active or idle. On the other hand, affinities measured *in vitro* with reconstituted vesicles or in heterologous systems cannot capture the effect of post-translational regulations that will arise in the original host organism.

Recently developed genetically-encoded, transport activity biosensors have been proposed as a way to overcome the difficulties associated with traditional characterization methods. These biosensors consist of fusions between full-length transporters and fluorescence proteins, where binding of the substrate to the transporter domain is translated into fluorescence changes (4, 5). These types of sensors are functional and can be used to replace the wild type transporter in plants, offering the opportunity to visualize substrate movement between cells and tissues (6).

Before transport activity biosensors become widely adopted in plant research, two critical limitations need to be addressed. First, the design of biosensors is laborious and requires multiple rounds of optimization. Given the large number of transporters present in plant species (1), progress in the field will depend on adopting techniques that will accelerate their generation. Second, it is still unclear how to reconcile the fluorescence changes displayed by transport activity biosensors with our molecular understanding of how transporters function. Here, we propose a design pipeline and a mathematical framework to address both of these concerns in the context of a new class of sugar transport activity biosensors, which we named SweetTracs. We constructed these sensors from *Arabidopsis thaliana* SWEET transporters, a seven-transmembrane domain family of mono- and disaccharide carriers. SWEETs are the most recently discovered family of transporters and the only known to facilitate the cellular release of sugars in plants (12).

## Results

### Generation of SweetTrac1

SweetTracs are intramolecular fusions between SWEETs and a circular-permuted superfolder Green Fluorescent Protein (cpsfGFP) (7). We created our biosensors by inserting cpsfGFP between the two pseudo-symmetric halves of *Arabidopsis* SWEET1 (Fig.1A). This approach was first demonstrated successful for AmTrac, a biosensor reporting ammonium transceptor activity (8). The insertion position was determined from a homology model of AtSWEET1 based on the structure of the rice SWEET2b (9). This position was confirmed using a yeast complementation assay to test for transport activity following the insertion of cpsfGFP at multiple locations between the third and fourth transmembrane domains (data not shown). Only chimeras that can still transport their substrate display a fluorescence change (8).

**Figure 1.**
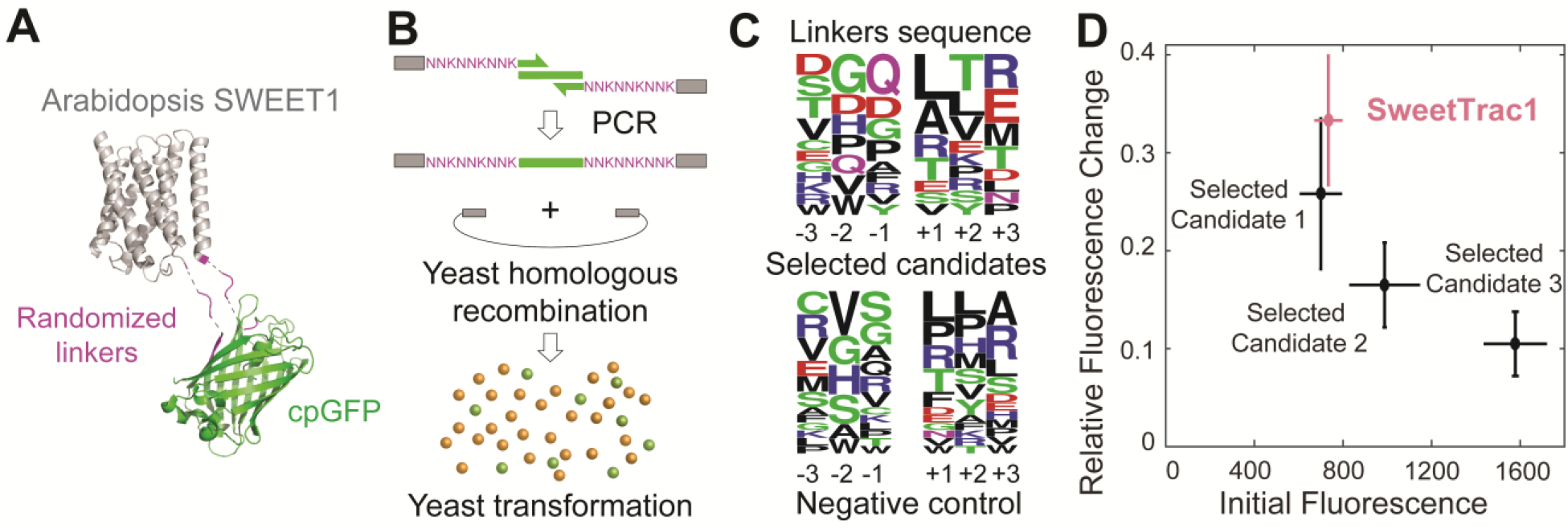
Generation of the SweetTtrac1 biomolecular sensor. (*A*) Molecular model of the SweetTrac1 sensor based on a homology model created from the structure of the rice SWEET2 (PDB ID 5CTG) and cpGFP (PDB ID 3EVP). (*B*) A pool of constructs with different linkers connecting SWEET1 and cpsfGFP were generated with mixed-base primers and isolated by FACS. (*C*) Amino acid distribution at each linker position for constructs that showed a change in fluorescence intensity in a screen with glucose. Weblogos were created from the amino acid frequency of 15 unique linker combinations. (*D*) Relative fluorescence change of the SweetTrac1 sensor (488 nm Exc, 514 nm Em), designed with the most conserved residues at each linker position, compared to the top three candidates identified during the screen.

The transporter and cpsfGFP in SweetTrac are connected by two linker peptides (Fig.1A). The composition of these linkers is known to be critical for biosensor performance. In AmTrac (8), for example, modifying the linkers varied the fluorescence intensity change resulting from ammonium addition. Unfortunately, the sequence space for linkers is vast, and optimization remains mostly empirical.

To accelerate the selection of linkers for SweetTrac, we took advantage of Fluorescence-Activated Cell Sorting (FACS) to screen a library of fusion proteins (Fig.1B-C). The library was generated by PCR amplification of the sequence of cpsfGFP using primers containing NNK degenerate codons to code for the linkers. The resulting DNA fragment was inserted by yeast homologous recombination into a vector containing the sequence of AtSWEET1 linearized at amino acid position 93 (Fig.1B). We screened 450,000 cells expressing biosensor variants in three separate experiments and isolated over 900 cells with the highest level of green fluorescence.

Cells isolated by FACS were regrown in liquid media and tested for fluorescence change with the addition of glucose, the substrate of AtSWEET1. We selected 44 outliers with the largest fluorescence increases for sequencing. We also chose a random set of 40 negative controls for comparison and observed apparent differences in amino acid composition (Fig.S1). We tested both 2 and 3 amino-acid-long linkers and found the latter to be the optimal length. We identified biosensor candidates with the same linker composition multiple times. There were 15 unique linker combinations amongst our 44 outliers. This result was unexpected, given the vast sequence space. We hypothesize that a combination of PCR amplification biases, low transformation efficiency, and the need to allow cells to recover and regrow following transformation may have reduced the diversity of our gene pool. Future improvements in gene synthesis and yeast transformation would increase the efficiency of our approach.

Sequence comparison of the linkers showed the predominance of particular amino acids at each position (Fig.1D). Furthermore, statistical coupling analysis suggests a lack of correlation between these amino acids (10), albeit a larger sample size may be necessary to corroborate this result. This led us to reason that optimal linkers may be obtained by selecting the predominant amino acid at each position. Indeed, we designed a variant with the predominance residues at each position (Fig.1E)—which we did not find in the screen—and it showed a larger response than the top three candidates identified during the screen. We named this designed variant SweetTrac1.

Furthermore, we were able to create a second biosensor from AtSWEET2, a vacuole sugar transporter that facilitates sugar storage in roots (11). We inserted the same cpsfGFP and linkers identified in our screen between the two pseudo-symmetric halves of AtSWEET2. The new biosensor, SweetTrac2, responded to the addition of glucose in a similar manner to SweetTrac1 (Fig.2A-D). This result suggests that our approach for designing biosensors could be generalized to other transporters. For instance, despite being closely related, AtSWEET1 and AtSWEET2 only share 44% sequence identity (Fig.S3). The ability to quantify the activity of transporters localized to intracellular membranes could be especially significant since they cannot be studied in whole cells with radiotracer uptake assays.

**Figure 2.**
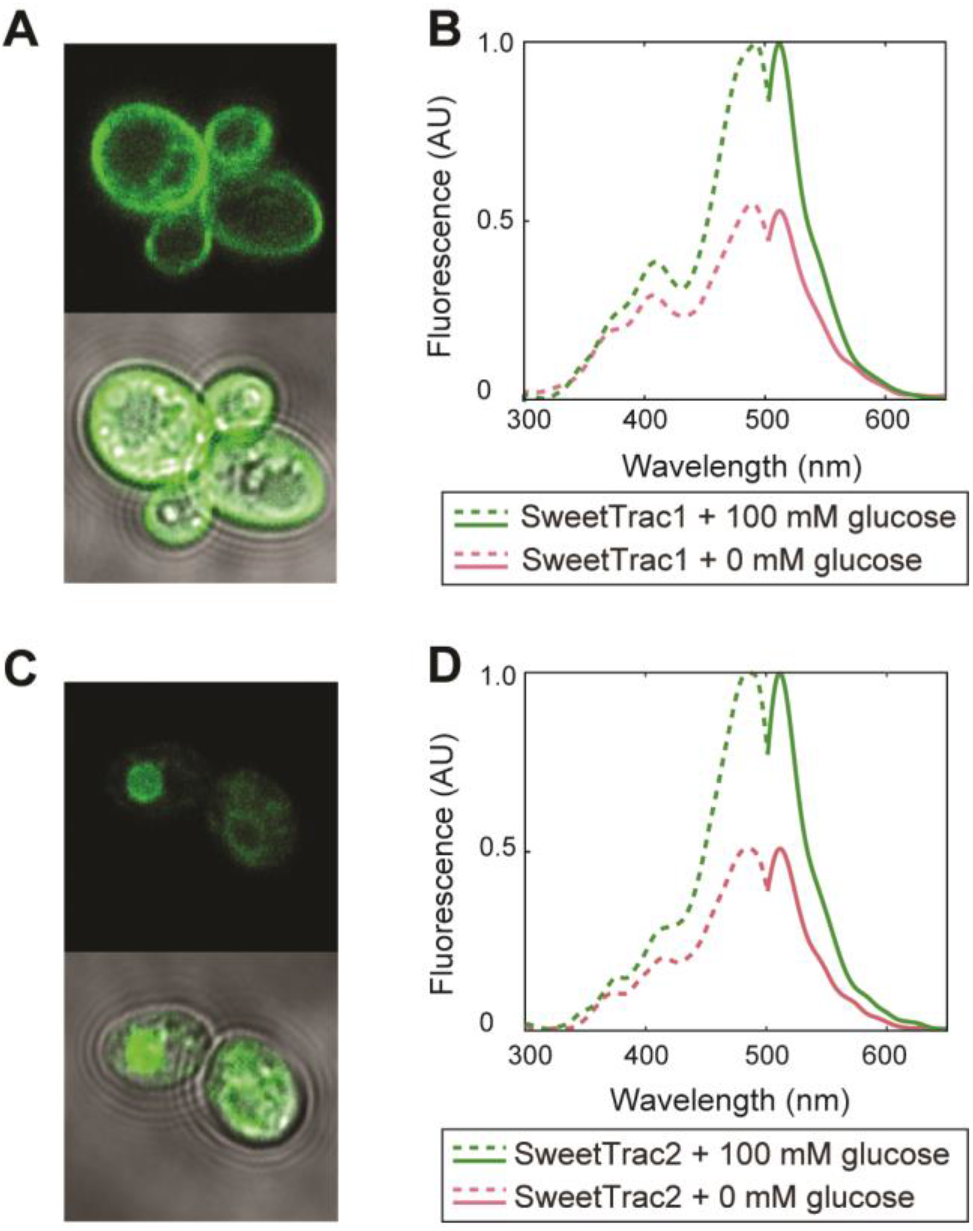
Characterization of SweetTrac sensors. (*A*) Localization of SweetTrac1 to the plasma membrane in yeast cells. (*B*) Normalized fluorescence excitation and emission spectra of SweetTrac1 (455 nm Exc, 530 nm Em). (*C*) Localization of SweetTrac2 to the tonoplast in yeast cells. Two focal planes (surface of a vacuole and cross-section) are shown. (*D*) Normalized fluorescence excitation and emission spectra of SweetTrac2 (455 nm Exc, 530 nm Em).

### Photophysical characterization of SweetTrac1 and SweetTrac2

Subcellular localization of SweetTrac in yeast cells mirrored that of the natural transporters *in planta* (11, 12). SweetTrac1 localized to the plasma membrane while SweetTrac2 localized to the vacuole (Fig.2A,C).

Spectra analysis of both SweetTracs revealed two excitation maxima—a major peak from the deprotonated chromophore at a wavelength of ~490 nm and a minor peak from the protonated chromophore at a wavelength of ~410 nm. A single emission maximum was observed at a wavelength of ~515 nm (Fig. 2B, D). The peak fluorescence intensity increases with glucose addition, and no shift in excitation and emission maxima was observed (Fig. 2B,D).

### SweetTrac1 reports glucose transport

AtSWEET1 had previously been shown to transport glucose (11, 12). To test if SweetTrac1 can transport glucose, we expressed SweetTrac1 in EBY4000, a yeast mutant lacking all endogenous hexose transporters, which prevent it from growing in media containing glucose as the sole carbon source (13). We found that the expression of SweetTrac1 in yeast rescued growth on liquid medium containing glucose, providing evidence that SweetTrac1 is a functional glucose transporter (Fig. 3A). Together, our data show that SweetTrac1 is a functional biosensor capable of reporting glucose transport.

**Figure 3.**
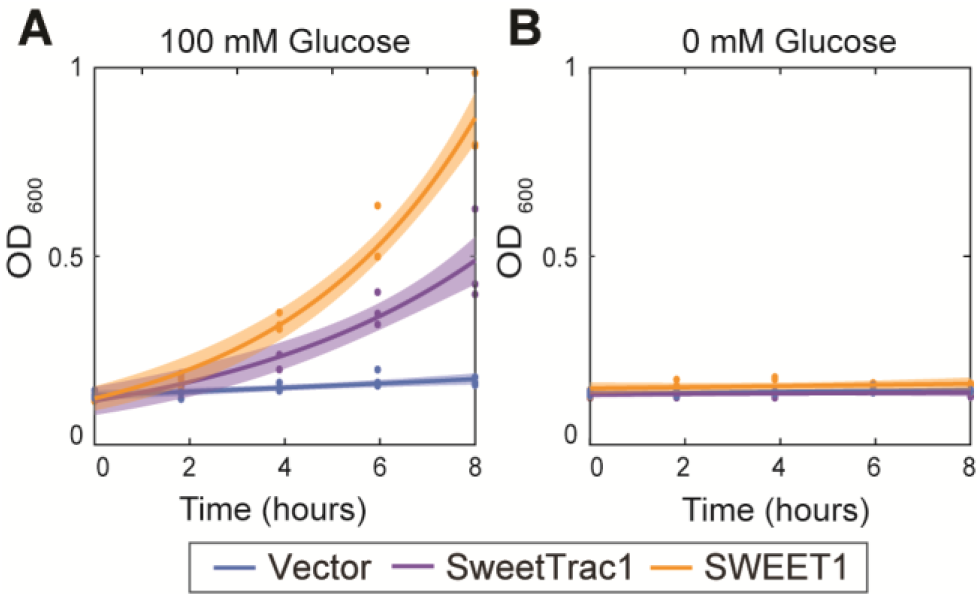
SweetTrac1 can transport glucose. (*A*) SweetTrac1 and SWEET1 can rescue the growth defect of the yeast strain EBY4000 in liquid media where glucose is the sole carbon source.

### Three-state uniporter model of SweetTrac1

The crystal structures of multiple bacterial homologs of SWEETs, the SemiSWEETs, show that these transporters adopt at least three states during a translocation cycle: outward-facing open, occluded, and inward-facing open (9, 14, 15). The identification of these states indicates that SWEETs follow an alternating access mechanism (15). SWEETs are also pH-independent and low-affinity transporters (1, 12). These characteristics are in alignment with a uniport mechanism or facilitated diffusion. Based on these observations, we propose a three-state, alternating access uniporter model for SweetTrac1 (Fig. 4A). In the model, *E_i_, E_o_*, and *ES* denote the inward-facing open, outward-facing open, and occluded bound state, respectively. The subscripts *i* and *o* indicate intra- and extracellular, respectively. In the case of vacuolar transporters, *i* and *o* denote intrafacial (cytosol) extrafacial (vacuole lumen), respectively. Six kinetic rate constants at which the transporter shifts between each configuration are denoted by *k*_1_-*k*_6_.

**Figure 4.**
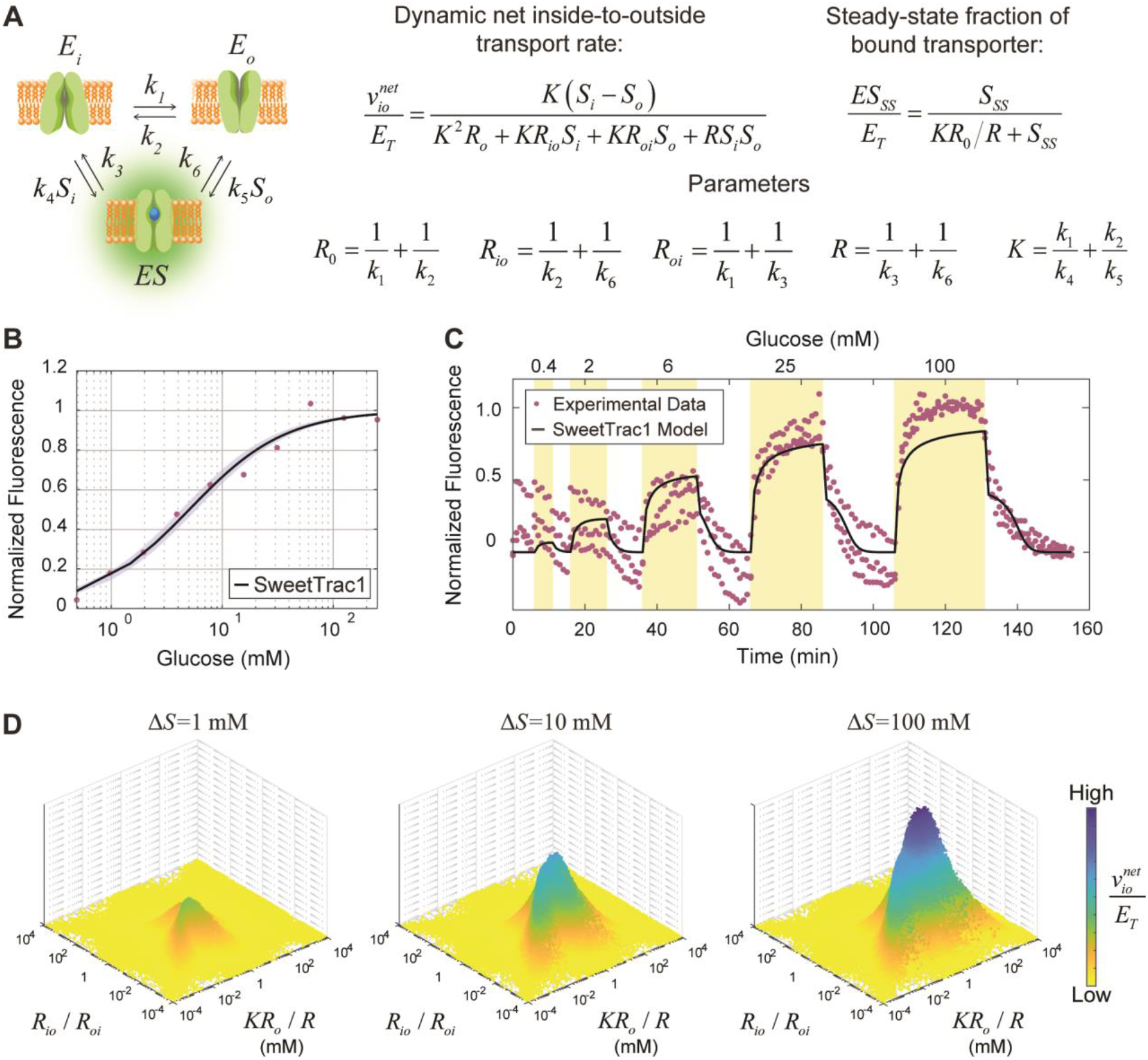
Kinetic model for uniporters and analysis of SweetTrac1. (*A*) Minimal, three-state model for different states of the transporters: Inside open (*E_i_*), outside open (*E_o_*), and substrate bound (*ES*). The net transporter rate and the steady-state fraction of bound transporter can be characterized by four resistance parameters (*R*_0_, *R_io_*, *R_oi_,* and *R*) and an intrinsic dissociation constant (*K*). At steady-state, the concentration of substrate (*S_ss_*) inside (*S_i_*) and outside (*S_o_*) of cells are equal. (*B*) Steady-state response of SweetTrac1 measured in microplate readers. Black line illustrates the model fit for *K*_*R*0_/*R* = 5.068±0.723 mM. (*C*) Dynamic response of SweetTrac1 measured in a commercial microfluidic device. The sensor showed a concentration-dependent increase in fluorescence when exposed to glucose pulses of different concentrations. Black line illustrate model fit for **K*_*R*0_/*R** = 5.068 mM and *R_io_/R_oi_* = 1.213±0.241. (*D*) Effect of different parameter values on the net transport rate at different concentration gradients (Δ*S*, where *S_o_* is set at 0.01 mM).

A net transport rate is derived from this model by making four major assumptions: (i) rates are determined by mass action kinetics, (ii) intermediate forms of the transporter (*E_i_, E_o_,* and *ES*) reach quasi-steady-state, (iii) each transition is in equilibrium with its reverse process (Principle of Detailed Balance), and (iv) measured fluorescence corresponds to the concentration of the bound state (*ES*). The basis for the first three assumptions is detailed in (16). The fourth assumption is based on the observation that the fluorescence of SweetTrac1 increases with glucose addition (Fig.2A). The *E_i_* and *E_o_* states of the sensor are then considered low fluorescence states, while the *ES* is a bright state.

The net transport rate can be described in terms of an intrinsic dissociation constant *K* and four resistances *R*, *R*_0_, *R_io_,* and *R_oi_* (Fig. 4A). The derivation for the rate equation and the molecular interpretation of each parameter is detailed in the SI Appendix following (16). Briefly, *R* and *R*_0_ are the overall resistances to a complete transport cycle by a bound and empty transporter, respectively. On the other hand, *R_io_* and *R_oi_* are the resistances in zero-trans experiments in the inside-to-outside and outside-to-inside directions, respectively (16).

### Model recapitulates the dynamics of SweetTrac1

The steady-state fraction of bound transporter can be described by a simple group of parameters, *KR*_0_/*R* (Fig.4A). This group represents the half-saturation concentration at steady-state when the intra- and extracellular concentrations of the substrate have equalized (*S_i_=S_o_*). Here, we will refer to this group as the *equilibrium exchange constant,* which can be used to describe the concentration-dependent response of our biosensors to sugars in microplate readers (Fig.4B). The equilibrium exchange constant for SweetTrac1 is 5.068±0.723 mM for glucose, meaning that half of the sensors will be in their bound state at this concentration. A similar analysis using SweetTrac2 estimated an equilibrium exchange constant of 4.441±0.985 mM for glucose (Fig.S2).

Our model correctly recapitulates the reversible and concentration-dependent behavior of SweetTrac1. Yeast cells expressing the biosensor were imaged in a fluorescence confocal microscope with the help of a commercially available microfluidic device. The device allowed the immobilization of the cells in the field of view and quick changes in the extracellular concentration of glucose (Fig.4C).

To estimate the intrinsic dissociation constant and resistances of SweetTrac1, we numerically solved the original system of four differential equations. The system of differential equations describes the intracellular accumulation of glucose and transporter fraction at each possible conformation (SI Appendix). We used this solution to fit the steady-state and dynamic fluorescence response of SweetTrac1 in the microfluidic device (Fig. 4B-C) to determine the rate constants *k*_1_-*k*_6_.

The parameters estimated in our model agree with experimentally established values. In a zero-trans influx experiment, the intracellular concentration of the substrate is zero, and the net transport rate equation reduces to a simpler form (*S_i_* = 0) that depends on the extracellular concentration of substrate and a group parameter *K*_*R*0_/*R_oi_* (SI Appendix). Here, we name this group the *zero-trans influx constant,* which is sometimes referred to as the Michaelis constant (*K*_m_) for uptake assays in literature. We found the value of the zero-trans influx constant for SweetTrac1 to be 3.8 ± 1.1 mM. This value is within the same order of magnitude than the 9.1 ± 1.2 mM determined for AtSWEET1 in influx assays of radiolabeled substrates (12). Differences in the values of these constants for the biosensor and the natural transporter suggest that the insertion of the cpsfGFP in AtSWEET1 may have affected its affinity. Nevertheless, SweetTrac1 is still expected to function within physiologically relevant concentrations *in planta.*

Our estimated parameters suggest that AtSWEET1 is a near symmetric transporter. Asymmetry is mathematically defined by the ratio of the resistance parameters *R_io_*/*R_oi_*. Uniporters are often kinetically asymmetric for a biological purpose (16). In red blood cells, for example, GLUT1 is highly asymmetric, with an *asymmetry ratio* of 18.75-37.5. This ratio indicates that GLUT1 greatly favors influx as opposed to efflux (17). It is suggested that its sided behavior allows glucose-exhausted cells to replenish themselves upon recirculation into a high glucose environment rapidly. Glucose release would be delayed when transiting low glucose environments (18). For SweetTrac1, we were able to estimate an asymmetry ratio of 1.213±0.241, indicating that AtSWEET1 is a near symmetric transporter that marginally favors influx (Fig. 4C), and thus would not buffer changes in intracellular sugar concentrations like GLUT1.

Furthermore, in the absence of a substrate, the model suggests that SweetTrac1 favors the inward-open state as opposed to the outward-open state (*k*_1_/*k*_2_=0.757). This ratio indicates that the inward-open state *E_i_* is roughly 1.3 times more stable than the outer-open state *E_o_*. Our observation is in alignment with reported molecular simulations of the bacterial LbSemiSWEET (14). This slower transition to the outward-open state has been suggested to slow and limit the overall transport rate. While we did find the transition from the inward-open to the outward-open state to be slower, we did not observe this transition to be particularly limiting. Instead, the asymmetry seems to arise from the breakdown of the loaded transporter. Our model shows the transporter unloading to the inner-state to be slower than the outer-state (*k*_3_/*k*_6_=3.41).

### Uniporter model reveals the existence of a theoretical optimum

It has been suggested that uniporters like SWEETs have an optimal affinity that parallels the physiological concentration of substrates at which they function (19). For efflux, when the transporter affinity is higher than the intracellular concentration of the substrate, the net efflux rate will be hindered by the lack of substrate. On the other extreme, when the affinity is considerably lower than the intracellular concentration, efflux could still be hindered. In this case, both the inward-facing and the outward-facing binding sites of the transporter would saturate with the substrate, the unidirectional inside-to-outside and outside-to-inside transport rate will have the same magnitude, and the net efflux rate will consequently approach zero.

We performed a computational analysis of our model to identify the conditions that lead to the optimal net transport rate. We explored a wide range of values of the rate constants *k*_1_-*k*_6_. We found that the maximum net transport rate depends on the interplay between the concentration gradient, the equilibrium exchange constant, and the asymmetry ratio (Fig. 4D). Maximum net transport occurs when the substrate concentration and the equilibrium exchange constant are within the same order of magnitude. Furthermore, a symmetric transporter could achieve higher net transport rates than an asymmetric one at any substrate concentration (Fig.4D). It is worth noting that we had a large number of parameters in our model and experimental noise in our parameter estimation. Despite our extensive search, we cannot exclude the existence of a different set of parameters that leads to a global minimum. More complex algorithms may be necessary to improve the parameter search.

Given the rate constants for our biosensors, we propose that SweetTracs, and implicitly SWEETs, operate optimally in the millimolar concentration range. This interpretation would suggest that low-affinity transporters like SWEETs have evolved to maximize net sugar transport at high sugar concentrations, such as those in plants, which can reach hundreds of millimoles (20).

## Discussion

In this work, we generated SweetTracs, new biosensors that report SWEET transport *in vivo,* and accompany it with a mathematical model to quantitatively explain their response. We used a unique method to study the structural transitions of membrane transporters without the need of radiolabeled or fluorescently-tagged substrates. Using SweetTrac1, we estimated the rate constants describing how SWEET1 shifts between conformations. To our knowledge, the estimation of the individual rate constants (not only the combined *K* and *R* parameters) has not been performed before.

Based on our analysis, we speculate that the physiological function of AtSWEET1 may be to equilibrate the intra- and extracellular concentrations of sugars rapidly. Given the low affinity and near symmetry demonstrated by our sensor (Fig.4D), we expect AtSWEET1 to allow high net influx or efflux rates *in planta.* This will rapidly equalize sugar concentrations on both sides of the membrane, unlike the “concentration buffering” attributed to GLUT1 in red blood cells (17, 18). While no mutant of AtSWEET1 has been reported yet, its maize homolog was recently shown to facilitate the influx of glucose into subsidiary cells, where glucose acts as a feedback inhibitor of stomatal opening (21). In this context, by allowing the intracellular concentration of glucose to equilibrate with the extracellular one quickly, ZmSWEET1 may help speed the response of the feedback mechanism. In general, SWEETs can be considered uniporters that, given their low affinity, are less likely to saturate at the high sugar concentrations common in plant tissues.

Sugar transporters like SWEETs play critical roles in many physiological processes, notably facilitating the efflux of sucrose out of the leaf cells—as the first step for phloem loading—and in pathogen infection (12, 22). This makes sugar transporters important targets for increasing crop yield and conferring pathogen resistance. Recently, genetic manipulation of transporters has been shown successfully in staple food crops such as potatoes and rice. In potato, overexpressing two metabolite transporters causes a dramatic increase in starch yield (23), while in rice, genomic editing promoters of SWEET genes conferred resistance to bacterial blight (24). Crop improvement could be made even faster with the help of computational models to select target genes (25). To realize such gains, we must first gather quantitative, real-time, and spatially-resolved information on metabolite concentrations and the activity of the cell machinery. Fluorescent biosensors can be invaluable tools for collecting such data non-destructively.

While our FACS-based approach to accelerate the development of biosensors was successful, some limitations are worth noting. When generating SweetTrac1, our screen for possible candidates was small compared with the theoretical number of possible variants (64 million variants for 6 amino acids). We can not rule out that upon further experimentation, we may find better linkers for our biosensors. In addition, the zero-trans efflux constant determined in our analysis of SweetTrac1 is higher than that determined for the wild-type SWEET1. The difference in estimated affinities may be attributed to the rigidity of the linkers and the cpsfGFP. This could also explain the decreased ability of SweetTrac1 to complement yeast growth in glucose as compared to wild-type SWEET1 (Fig. 3A). Further improvements to our strategy, such as the use of high-throughput sequencing of pooled candidates to increase the number of linkers analyzed, may help optimize their composition. Nevertheless, our positive results do suggest that exhaustive coverage of the initial gene library may not be necessary.

We are in the midst of a high-throughput revolution, where plant genomes are becoming readily available. Approximately a tenth of these genomes encode transport proteins (26). As we continue to identify transporters in larger numbers, we will need new functional characterization methods that match the speed of current sequencing techniques. We will also need ways to study transporters in their physiological context, in association with their regulatory proteins and metabolic pathways. Biosensors could be important tools to facilitate such studies (6, 27), and our work demonstrates their potential for collecting quantitative data. Furthermore, sensors like our SweetTracs can help answer system-level questions on substrate competition, concentration robustness, and many more.

## Materials and Methods

### DNA constructs

The *Arabidopsis* SWEET1 (At1g24300) and SWEET2 (At3g14770) CDS were used as the basis for generating the SweetTrac sensors. All sequences were cloned into the entry vector pDRf1-GW for yeast expression (28) through Gateway cloning (Invitrogen).

For the construction of the SweetTrac1 library, an XbaI restriction site was inserted into position 93 of the AtSWEET1 protein. The modified pDRf1-SWEET1 plasmid was then linearized with XbaI and gel purified. Mixed-base primers, where the linkers coding sequences were randomized using NNK degenerate codons, were used to PCR amplify the sequence of cpsfGFP. NNK encodes for 32 codons: 20 amino acids and 1 stop codon. The linearized pDRf1-SWEET1 vector and cpsfGFP fragments were ligated via homologous recombination in *S. cerevisiae*.

The final SweetTrac1 and SweetTrac2 sensors contain the coding sequence of cpsfGFP, flanked by the left and right amino acid linkers DGQ and LTR, respectively. The linkers and cpsfGFP were inserted into amino acid position 93 in SWEET1 and 102 in SWEET2.

### Yeast transformation and sorting

The SWEET1-cpsfGFP fusion library was expressed in *S. cerevisiae* using the lithium acetate method (29) and plated on solid Synthetic Minimal (SD) medium supplemented with 2% agar, 2% maltose, and auxotrophic requirements. After 24 hours, yeast cells were recovered by rising agar plates with liquid SD medium supplemented with 2% maltose and auxotrophic requirements. Cells were sorted in a FACSAria II (BD Biosciences). Fluorescence was excited using a 488 nm laser line, and emission collected with a 530/30 nm filter set. A total of ~450,000 cells were screened over three independent experiments, and over 900 cells (0.2%) with the highest level of green fluorescence were isolated for further testing.

To identify chimeras that exhibit a fluorescence change with the addition of substrate, colonies were regrown overnight in 96-well blocks with YP (Yeast extract-Peptone) medium supplemented with 2% maltose, washed thrice with deionized water, and resuspended in SD medium supplemented only with auxotrophic requirements. The fluorescence intensity of each culture (at an excitation wavelength of 488 nm and an emission wavelength of 514 nm) was recorded before and after the addition of 100 mM of glucose using a Tecan M1000 Pro plate reader. Plasmids from 44 colonies that exhibited a statistically significant change in fluorescence intensity with the addition of glucose were recovered and sequenced. An additional 40 colonies were randomly selected from the remaining pool and sequenced for comparison (Fig.S1). The frequency of different amino acids in unique linkers is illustrated with web-based application WebLogo (30).

### Fluorimetric analyses

The SweetTracs were expressed in the yeast strain 23344c (*MATα ura3-52),* and the fluorescence was recorded in a Tecan Spark plate reader in clear, 96-well microplates. Yeast cells expressing the empty pDRf1 vector were used for background subtraction.

To measure the excitation and emission spectra, yeast cultures were washed twice with deionized water and suspended in liquid SD medium to an OD_600_ of 0.5. Excitation and emission spectra were recorded at an emission wavelength of 530 nm and an excitation wavelength of 455 nm, respectively.

To measure the steady-state fluorescence response of SweetTrac1 and SweetTrac2, cultures were washed twice and suspended in liquid SD medium to an OD_600_ of 0.9. Fluorescence was measured at an emission wavelength of 488 nm and an excitation wavelength of 514 nm.

### Yeast growth assay

The yeast strain EBY4000 (*hxt1-17D::loxP gal2D::loxP stl1D::loxP agt1D::loxP ydl247wD::loxP yjr160cD::loxP)* was used for yeast complementation assays (13). Single colonies were inoculated in liquid SD medium supplemented with 2% glucose and grown at 30 °C until an OD_600_ of 0.3-0.5. The yeast cells were washed twice with deionized water, then resuspended with liquid SD medium either supplemented with or without 100 mM of glucose to an OD_600_ of 0.1. The cultures were incubated in 96-well blocks (Zymo) at 30 °C with agitation for a total of 8 hours, taking OD_600_ readings at 2 hours intervals.

### Yeast imaging conditions

A Zeiss LSM 700 with a 63x oil, 1.4 NA objective laser scanning confocal microscope was used to capture the subcellular localization of SweetTrac1 and SweetTrac2 (Fig.2A,C). Fluorescence was excited using a 488 nm laser line, and emission collected with a BP 525/50 nm filter set (Carl Zeiss Microscopy).

A Yokogawa CSU-X1 (Mitaka) spinning disk confocal microscope equipped with a Photometrics Evolve EMCCD camera was used to collect the dynamics of the SweetTrac1 sensor (Fig.4C). Yeast cells were trapped as a monolayer in a microfluidic perfusion system (CellASIC) and exposed to different concentrations of glucose. Sorbitol was used as a counter osmolyte in the wash buffer. Fluorescence was excited using a 488 nm laser line, and emission was collected with a 525/50 nm filter set (Semrock).

### Computational and statistical analyses

All computational analyses and plots were generated using MATLAB (MathWorks).

Yeast growth was fit to an exponential growth curve OD_600_ = *ae_kt_*, where *a* is the initial OD_600_ and *k* is the growth constant. SweetTrac1 response was fit to the steady-state fraction of the bound transporter. Both expressions were fit to experimental data using MATLAB’s curve fitting toolbox (Fig.4A). For both fits, we used a nonlinear least-squares fitting procedure with a trust-region algorithm. Parameters and plots are reported with 95% confidence intervals.

The kinetic model for uniporters was used to analyze the effect of rate parameters *k_i_* on the net transport rate of glucose *v_io_.* For the analysis, 1.8×10^6^ different combinations of parameter values ranging from 10^-3^ to 10^4^ were evaluated at varying glucose concentration gradients. For details on the parametric fitting of SweetTrac1’s dynamic response data, see SI Appendix.

## Acknowledgments

We thank Cindy Ast and Mark P. Styczynski for helpful discussions and comments on the manuscript. This work was supported by National Science Foundation grants 1401855 and 1942722 to LSC. Contributions by WBF were supported by Deutsche Forschungsgemeinschaft (DFG, German Research Foundation) under Germany’s Excellence Strategy – EXC-2048/1 – project ID 390686111, as well as the Alexander von Humboldt Professorship.

## Notes

### Competing Interest Statement

The authors have declared no competing interest.

