## Supplementary material for "Quantitative analysis of transporter activity biosensors": SI Appendix

* Lily S. Cheung

**This PDF file includes:**

Supplementary text
Figures S1-S3

**Derivation of the net export rate and the steady-state fraction of bound transporter for a three-state model.**

The rate of change of each species is given by:

$$\frac{dS_i}{dt} = k_3 ES - k_4 E_i S_i \tag{1}$$

$$\frac{dE_i}{dt} = -k_1 E_i + k_2 E_o + k_3 ES - k_4 E_i S_i \tag{2}$$

$$\frac{dE_o}{dt} = k_1 E_i - k_2 E_o - k_5 E_o S_o + k_6 ES \tag{3}$$

$$\frac{dES}{dt} = -k_3 ES + k_4 E_i S_i + k_5 E_o S_o - k_6 ES \tag{4}$$

From the principle of detailed balance, the forward rates and the reverse rates are equal at equilibrium:

$$k_1 k_3 k_5 = k_2 k_4 k_6 \tag{5}$$

The total amount of transporter is conserved:

$$E_T = E_o + E_i + ES \tag{6}$$

Applying the quasi-steady-state assumption:

$$\frac{dE_o}{dt} = \frac{dE_i}{dt} = \frac{dES}{dt} = 0 \tag{7}$$

The fraction of the transporter at the three different states can be calculated from equations (2), (3) and (6)

$$\begin{bmatrix} -k_1 - k_4 S_i & k_2 & k_3 \\ k_1 & -k_2 - k_5 S_o & k_6 \\ 1 & 1 & 1 \end{bmatrix} \begin{bmatrix} E_i \\ E_o \\ ES \end{bmatrix} = \begin{bmatrix} 0 \\ 0 \\ E_T \end{bmatrix}$$

Solving the system of algebraic equations, we obtain:

$$\frac{E_i}{E_T} = \frac{k_2 k_3 + k_2 k_6 + k_3 k_5 S_o}{(k_3 + k_6)(k_1 + k_2) + k_4 (k_2 + k_6) S_i + k_5 (k_1 + k_3) S_o + k_4 k_5 S_i S_o} \tag{8}$$

$$\frac{E_o}{E_T}=\frac{k_1k_3+k_1k_6+k_4k_6S_i}{(k_3+k_6)(k_1+k_2)+k_4(k_2+k_6)S_i+k_5(k_1+k_3)S_o+k_4k_5S_iS_o} \quad (9)$$

$$\frac{ES}{E_T}=\frac{k_1k_5S_o+k_2k_4S_i+k_4k_5S_iS_o}{(k_3+k_6)(k_1+k_2)+k_4(k_2+k_6)S_i+k_5(k_1+k_3)S_o+k_4k_5S_iS_o} \quad (10)$$

The unidirectional export rate of substrate is equal to the product of the rate of formation of the complex and the fraction of the complex that releases the substrate to the extracellular side:

$$v_{io}=k_4E_iS_i\left(\frac{k_6}{k_3+k_6}\right) \quad (11)$$

$$\frac{v_{io}}{E_T}=\frac{\left[\left(\frac{k_2}{k_3k_5}\right)(k_3+k_6)+S_o\right]S_i}{\frac{(k_3+k_6)^2(k_1+k_2)}{k_3k_4k_5k_6}+\frac{(k_3+k_6)(k_1+k_3)}{k_3k_4k_6}S_o+\frac{(k_3+k_6)(k_2+k_6)}{k_3k_5k_6}S_i+\frac{(k_3+k_6)}{k_3k_6}S_iS_o}$$

From the first term in the numerator, we obtain the intrinsic affinity constant $K$:

$$\left(\frac{k_2}{k_3k_5}\right)(k_3+k_6)=\frac{k_2}{k_5}+\frac{k_2k_6}{k_3k_5}=\frac{k_2}{k_5}+\frac{k_1}{k_4}=K$$

$K$ represents the half-saturation concentration when there is no substrate at either sides of the membrane. Its reciprocal is the intrinsic affinity that is unperturbed by the presence of substrate.

From the first term in the denominator, we obtain the resistance parameter $R_0$:

$$\frac{(k_3+k_6)^2(k_1+k_2)}{k_3k_4k_5k_6}=\left(\frac{k_1k_3}{k_4k_6}+\frac{k_2k_6}{k_3k_5}\right)^2\left(\frac{k_1+k_2}{k_1k_2}\right)=K^2R_0 \text{ , where } \frac{1}{k_1}+\frac{1}{k_2}=R_0$$

$R_0$ is the overall resistance to the complete transport cycle by the unloaded transporter.

From the second term in the denominator, we obtain the resistance parameter $R_{oi}$:

$$\frac{(k_3+k_6)(k_1+k_3)}{k_3k_4k_6}=\left(\frac{k_1k_3}{k_4k_6}+\frac{k_2k_6}{k_3k_5}\right)\left(\frac{k_1+k_3}{k_1k_3}\right)=KR_{oi} \text{ , where } \frac{1}{k_1}+\frac{1}{k_3}=R_{oi}$$

$R_{oi}$ is the resistance to the transport cycle in a situation where the extracellular side of the membrane is limitingly high (zero-trans).

From the third term in the denominator, we obtain the resistance parameter $R_{io}$:

$$\frac{(k_3+k_6)(k_2+k_6)}{k_3k_5k_6}=\left(\frac{k_1k_3}{k_4k_6}+\frac{k_2k_6}{k_3k_5}\right)\left(\frac{k_2+k_6}{k_2k_6}\right)=KR_{io}$$ , where $\frac{1}{k_2}+\frac{1}{k_6}=R_{io}$

$R_{io}$ is the resistance to the transport cycle in a situation where the intracellular side of the membrane is limitingly high (zero-trans).

From the last term in the denominator, we obtain the resistance parameter $R$:

$$\frac{k_3+k_6}{k_3k_6}=\frac{1}{k_3}+\frac{1}{k_6}=R$$

$R$ is the overall resistance to the complete transport cycle by the loaded transporter.

Thus, unidirectional export rate of the substrate becomes:

$$\frac{v_{io}}{E_T}=\frac{(K+S_o)S_i}{K^2R_o+KR_{io}S_i+KR_{oi}S_o+RS_iS_o} \quad (12)$$

The net unidirectional export rate of substrate becomes:

$$\frac{v_{io}^{net}}{E_T}=\frac{K(S_i-S_o)}{K^2R_o+KR_{io}S_i+KR_{oi}S_o+RS_iS_o} \quad (13)$$

and the net unidirectional import rate of substrate becomes:

$$\frac{v_{oi}^{net}}{E_T}=\frac{K(S_o-S_i)}{K^2R_o+KR_{io}S_i+KR_{oi}S_o+RS_iS_o} \quad (14)$$

The fraction of bound transporter can be derived by substituting the definition of the intrinsic affinity constant and resistance parameters into equation (10):

$$\frac{ES}{E_T}=\frac{\left(\frac{k_3+k_6}{k_3k_6}\right)\frac{k_1}{k_4}S_o+\left(\frac{k_3+k_6}{k_3k_6}\right)\frac{k_2}{k_5}S_i+\left(\frac{k_3+k_6}{k_3k_6}\right)S_iS_o}{K^2R_o+KR_{io}S_i+KR_{oi}S_o+RS_iS_o}$$

At steady state, $S_i=S_o=S_{SS}$ ,

$$\frac{ES_{SS}}{E_T}=\frac{\left(\frac{k_1}{k_4}+\frac{k_2}{k_5}\right)RS_{SS}+RS_{SS}^2}{K^2R_o+K(R_{io}+R_{oi})S_{SS}+RS_{SS}^2}$$

and noticing that $R_{io} + R_{oi} = R_0 + R$, we obtain the expression:

$$\frac{ES_{SS}}{E_T} = \frac{S_{SS}}{\frac{KR_o}{R} + S_{SS}} \qquad (13)$$

The fraction of the transporter at steady-state is characterized by a single parameter, $KR_0/R$, which we call the equilibrium exchange constant.

**Interpretation of glucose uptake data in terms of uniporter model parameters.**

For a zero-trans experiment measuring efflux, $S_o$ is set to zero. Equation (13) reduces to:

$$\frac{v_{io}^{net}}{E_T} = \frac{\left(\frac{1}{R_{io}}\right) S_i}{\left(\frac{KR_0}{R_{io}}\right) + S_i}$$

We call the group $KR_0/R_{io}$ the *zero-trans efflux constant*, and it can in interpreted as the half-saturation concentration, when the intracellular concentration of substrate is limitingly high compared to the extracellular concentration.

Similarly, for a zero-trans experiment measuring uptake, $S_i$ is set to zero. Equation (14) reduces to:

$$\frac{v_{oi}^{net}}{E_T} = \frac{\left(\frac{1}{R_{oi}}\right) S_o}{\left(\frac{KR_0}{R_{oi}}\right) + S_o}$$

Where we refer to the group $KR_0/R_{ot}$ as the *zero-trans influx constant*, and it can in interpreted as the half-saturation concentration, when the extracellular concentration of substrate is limitingly high compared to the intracellular concentration.

**Parametric fitting of SweetTrac1's dynamic response data.**

The three-state uniporter model was used for parametric fitting of SweetTrac1's steady state and dynamic response data (Fig. 5C). The difference between the model and experimental data was set as a nonlinear minimization of least-squares problem. The system of ordinary differential equations describing the state dynamics of the model (1-4) were used.

Only parameters $k_1$, $k_2$, $k_3$, $k_6$, were estimated. $k_4$ and $k_5$ were calculated using the equilibrium rate constraint (5) and the equilibrium exchange constant (5.068 mM) determined from experiments in microplate readers (Fig. 5B).

95% confidence intervals for rate parameter estimates were obtained using MATLAB's nlpredci function from the Statistics Toolbox.

### Supplemental Figures

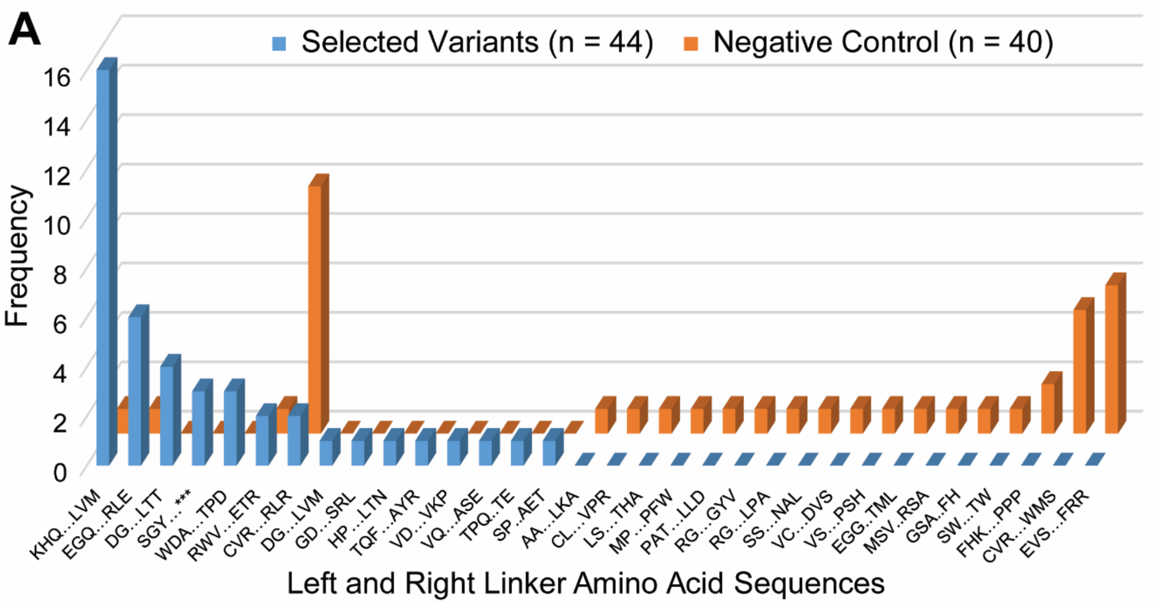


**Figure S1. Linker composition of biosensor candidates.** *(A)* The frequency of amino acid sequences for left and right linkers shown.

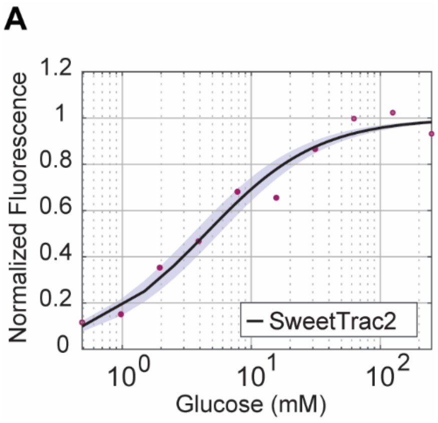


**Figure S2. Kinetic analysis of the SweetTrac2 sensor.** *(A)* Steady-state response of SweetTrac2 measured in microplate readers. Black line illustrates model fit for $KR_0/R$ = 4.441±0.985 mM.

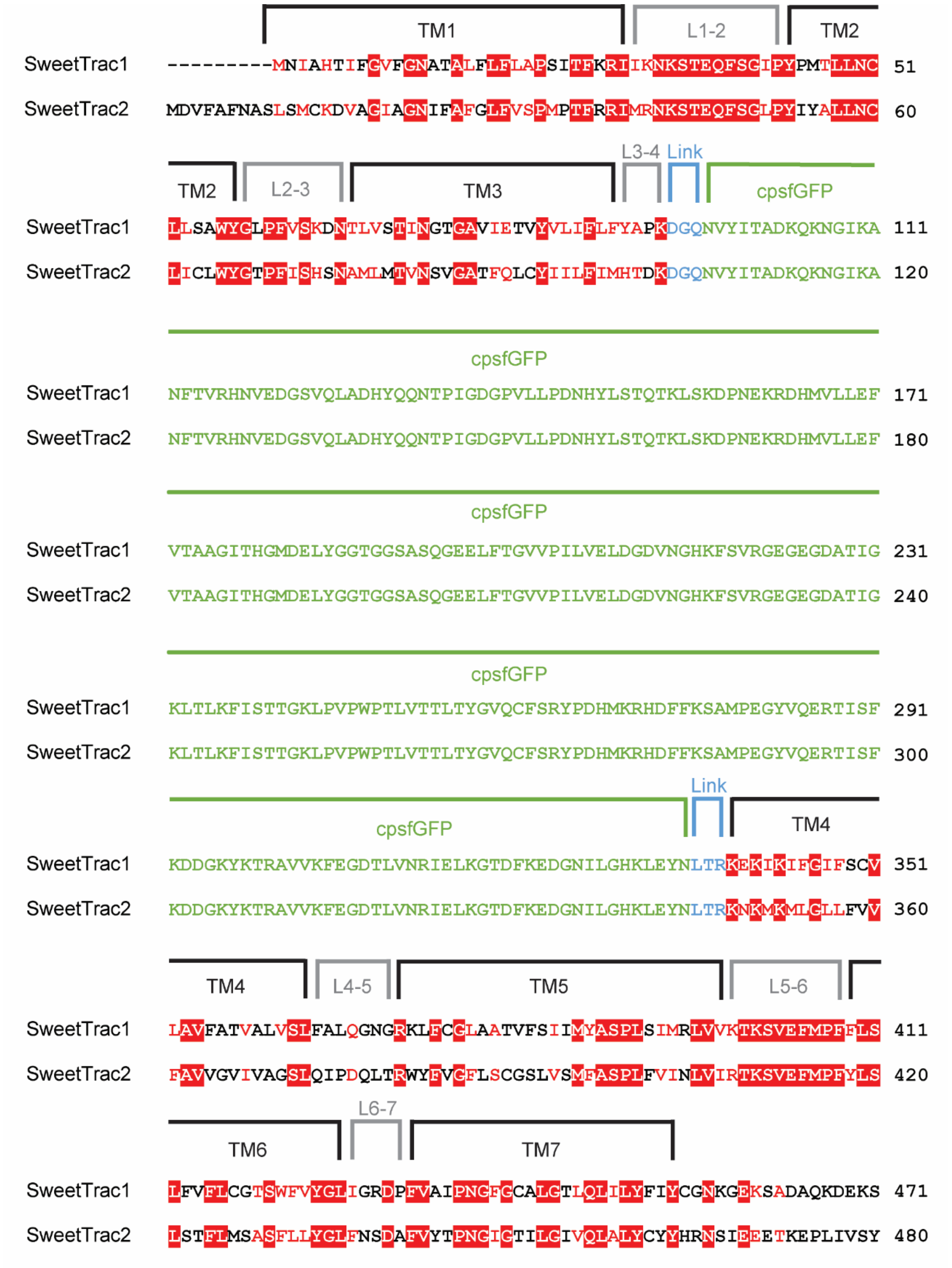


**Figure S3. Sequence alignment of SweetTrac1 and SweetTrac2.** *(A)* SweetTrac1 and SweetTrac2 sequences were aligned using Clustal Omega. The secondary structure marked above the sequences were based on OsSWEET2b.
